# Genotype of Human Papilloma virus in Male Genital Warts In Korean Men and Review of Literature

**DOI:** 10.1101/2021.07.19.453017

**Authors:** Woochul Moon, Jungho Jo, Jinhan Yoon, Korean Male HPV Study Group, Jung Joo Moon

## Abstract

**Purpose:** Genital warts are one of the most common sexually transmitted infections and are known to develop due to human papillomavirus (HPV) infection, especially HPV types 6 and 11. However, their prevalence and subtypes in male genital warts remains poorly defined. HPV vaccine is administered to men in part to prevent anogenital warts and it is important to investigate their expected impact in male anogenital warts.

**Materials and Methods:** We have herein conducted a multicenter, prospective study to analyze HPV type distribution in genital warts of 1000 Korean men by using DNA microarray that can detect 40 types of genital HPV.

**Results:** 1000 out of 1015 genital warts showed HPV DNA. Out of 1000 HPV-positive samples, 18.8% showed mixed infection and 81.2% showed single infection. Of 18 high-risk (16.2%) and 14 low-risk (94.3%) HPV types detected, the most common type of HPV types were HPV6 (59.5%), followed by HPV11 (24.3%), HPV16 (5.8%), HPV91 (5.3%), HPV40 (3.3%). 85.9% showed the 9 HPV types covered by the vaccine. Sixteen of the 200 HPV specimens submitted for sequencing showed discrepant results compared to the DNA sequencing.

**Conclusions:** Male genital warts predominantly show low-risk type HPV (HPV 6 and 11). However, high-risk HPV is not uncommon and the role of high-risk HPV in genital warts may be considered. The Gardasil 9 HPV vaccine is expected to provide protection against about >80% of male genital warts. Further HPV typing studies in male genital warts are necessary in other races and geographical areas to define the role and management of high-risk type HPV in male genital warts.

## INTRODUCTION

Human papillomaviruses (HPVs) are a group of small double-stranded DNA viruses that infect the human epithelium, causing hyperproliferation, and one of the most common sexually transmitted infections worldwide.^1^ There are over 200 types of HPV, of which approximately 45 types infect epithelial and mucosal lining of the anogenital area, which is called genital or anogenital HPV.^2^ HPV types are organized into five major genera: alpha, beta, gamma, mu, and nu, and HPV infections are divided into cutaneous and mucosal HPV.^2^ Cutaneous HPV causes common warts and mucosal HPV causes anogenital infections and lesions. Mucosal HPV induces a variety of external genital lesions, including warts (condylomata acuminata), precancerous lesions, and cancer, and is classified as high-risk or low-risk depending on its oncogenic potential.^2^ Nearly 100% of uterine cervix cancer, 36%–40 % vulvar cancer, close to 90 % of vaginal cancers in females and 80%–85% of anal cancers and close to 50% of penile cancers in males, and 20%–30% of head and neck cancers develop secondary to HPV infection.^3^ Cervical cancer is the second most common cancer affecting women.^1^ With the appreciation of the role of HPV in cervical carcinogenesis, HPV detection and genotyping has become a standard screening tool for cervical cancer in combination with cytology studies. ^4^

Quadrivalent HPV vaccine (Types 6, 11, 16, and 18; Gardasil; Merck, Sharp & Dohme Corp) was approved by the FDA (U.S. Food and Drug Administration) on June 8, 2006 for females 9-26 years of age to protect against cervical, vulvar and vaginal cancers and genital warts.^5,6.7^ Later, it was approved in both men and women 9 through 26 years of age for the prevention of genital warts and anal cancers.^6^ On December 10, 2014, the FDA approved Gardasil 9, which covers the same four HPV types as Gardasil, as well as an additional five HPV types (6, 11, 16, 18, 31, 33, 45, 52, and 58), for use in males and females aged 9 through 26 years and eventually quadrivalent Gardasil vaccine was discontinued.^5,7^ On October 5, 2018, FDA expanded the approved use of the Gardasil 9 vaccine to include women and men aged 27 through 45 years.^7^ HPV vaccination prior to becoming infected with the HPV types covered by the vaccine has the potential to prevent more than 90 percent of effected cancers, or 31,200 cases every year, from ever developing.^7^

Risk factors associated with persistence of HPV infection include older age, cigarette smoking or other tobacco use, immunocompromised status (including HIV), nutritional deficiencies, non-use of condoms, presence of other STDs, oral contraceptive use, uncircumcised status among men, and human leukocyte antigen (HLA) polymorphisms. ^9^

Although less appreciated, the burden of HPV in men is also significant and needs to be evaluated. HPV induces anal, penile, and head and neck cancers in men. ^4,10,12–13^ Most studies examining the role of HPV in the development of male external genital lesions (EGLs) have identified mucosal HPV.^10–13^ High- and low-risk HPV types are found in 15.6% and 73.2% of EGLs, HPV 6 or 11 in condylomas, HPV 16 in PEIN (penile intraepithelial neoplasia) I or II lesion and 1 PeIN III lesion positive for HPV 6 only in a study by DJ Ingles et al., while 70% and 100% of PEINs are HPV-positive, and 40 %–50% of invasive penile cancers are HPV positive in a study by Dillner et al.^11^

The major HPV-related diseases in men are genital warts. The majority of genital warts develop due to infection by low-risk HPVs.^12–14^ While HPV-related condyloma is considered a benign lesion, the substantial economic and psychosocial burden of this clinical manifestation of infection cannot be overlooked. HPV type 6 and 11 have been reported to cause more than 90% of genital warts. ^12–14^

However, contrary to the common belief that HPV 6 and HPV 11 induce almost all genital warts in males, and that vaccines targeting HPV 6 and HPV11 will provide protection against most of them, data on genotyping information of male genital warts are scarce. ^4, 16–18^ There have been suggestions that at least some genital warts, in addition to HPV6 and HPV11, also contained co-infection with high-risk HPVs.^17, 18^ Epidemiological studies indicate that the 9vHPV vaccine could prevent approximately 90% of cervical cancers, 70 %–85% of high-grade cervical dysplasia (precancers), 85 %–95% of HPV-related vulvar, vaginal, and anal cancers, and 90% of genital warts in women.^19^ Therefore, to review the effect of Gadasil 9 in the male population, it is pertinent to identify the prevalence of these nine HPV types in anogenital warts in men. Moreover, monitoring the impact of vaccination on HPV infection and disease in men raises challenges such as the long time frame until cancer outcomes and complexity of factors that need consideration (different policies, health system outcomes, and biological outcomes), as well as the fact that genital specimens suitable for monitoring HPV prevalence are not routinely collected for other diagnostic or screening purposes in males ^20^.

GG HPV DNA microarray (HPV40 DNA chip) is an oligonucleotide microarray that can detect 40 types of genital HPV in an accurate, high-throughput, and cost-effective way.^21^ We herein have carried out a large-scale genotyping study in Korean men with genital warts by using HPV DNA chip. The purpose of the current study was to identify the precise genotyping information of HPV in male genital warts, specifically to investigate whether genital warts contain high-risk HPVs, how much proportion of genital warts contain low-and high-risk HPVs, and to predict the potential efficacy of currently available HPV vaccines in the protection of male genital warts.

## MATERIALS AND METHODS

We investigated the genomic DNA of HPV from fresh tissues of pathologically diagnosed genital warts from 1015 Korean adult men using the GG HPV DNA chip. The GG HPV DNA chip (Goodgene Inc., Seoul, Korea) has multiple oligonucleotide probes for 40 types of genital HPV and human beta globin genes and identified 40 HPV types (HPV 6, 11, 16, 18, 26, 30–35, 39, 40, 42–45, 51–56, 58, 59, 61, 62, 66–70, 72, 73, 81–84, 90, and 91). The HPV DNA chip has been licensed by the Korean FDA for genotyping of genital HPV and screening of precancerous lesions and cancer of the uterine cervix. It can detect 10–100 copies of HPV per sample. Genomic DNA extraction, amplification, labeling, hybridization, and analysis were performed according to the manufacturer’s instructions. Briefly, genomic DNA was extracted using the LaboPassTM Tissue mini prep. Kit (Cosmo Genetech Products, Seoul, Korea). The primers chosen were the L1 gene: primers Ll and L3. PCR with the primers L1 and L3 amplified approximately 200-base pair DNA fragments of all genotypes of HPV. A mixture of 10 mL of HPV DNA-amplified product and 10 mL beta-globin-amplified products were denatured by heating at 95 °C for two min, followed by cooling for 3 min on ice. The samples were mixed with 65 mL of hybridization buffer (Goodgene, Seoul, Korea) and placed on the HPV DNA chip. The HPV DNA chip was incubated at 50 °C for 30 min. The HPV DNA chip was washed twice with 36 SSPE for 2 min and 16 SSPE for 2 min. This led to the formation of visible spots on the array surface, which were then scanned, measured, and analyzed using a dedicated reader and software after the chip was dried (Molecular MDC Genepix 4000A Scanner, Molecular Devices Inc., California, USA). To confirm and validate the results of the HPV DNA chip analysis, they were comparatively analyzed by DNA sequencing assay in selected cases.

### Patient recruitment and sample collection

With the appreciation of the importance of HPV in men, a group of urologists in Korea and molecular scientists established a “Korean Male HPV (KM-HPV) Study Group” in 2008, which aimed to study the prevalence and genotype of HPV infection and guide HPV vaccination in men. This study was carried out in this group. A total of 1050 patients were enrolled in the present study from 30 urologic clinics in Seoul and Busan, South Korea, over a 2-year period. Patients with a clinical diagnosis of genital warts were invited to participate in this study. In accordance with Korean regulations, because all patients presented through private urologic clinics and voluntarily signed informed consent, no ethics committee approval was necessary. This study complied with the Declaration of Helsinki, and written informed consent was obtained from each patient. Fresh tissue fragments of genital lesions consistent with exophytic condylomata acuminata were removed by excision biopsy in a strictly sterile manner at the time of operative treatment. Then, they were immediately immersed in 3 ml of BD Universal Viral Transport Medium (BD Diagnostics, Franklin Lakes, NJ, USA) and sent to the laboratory and stored at 4^0^ °C until processing. Slides of the specimen were made, stained with hematoxylin and eosin, and evaluated by pathologists for the presence of inflammatory, infectious, preneoplastic, or neoplastic conditions. Samples with coexisting neoplastic conditions, such as EIN or cancer, were excluded from this study.

### HPV detection and genotyping assay

#### DNA extraction and PCR

After receiving the samples, tissue fragments were fixed in formalin and embedded in paraffin. DNA was extracted from formalin-fixed, paraffin-embedded specimens using the QIAamp DNA FFPE Tissue Kit (Qiagen) according to the manufacturer’s protocol. The sample DNA was obtained from tissue containing both epithelium and stroma, so the concentration of HPV in infected keratinocytes is likely to be higher.^35^ Genotyping was performed to detect HPV DNA using the GG HPV DNA chip, which detects 40 HPV genotypes classified as high- or low-risk, depending on its association with the development of carcinoma (high-risk types: 16, 18, 26, 31, 33, 35, 39, 45, 51, 52, 53, 56, 58, 59, 66, 67, 68, 69, 73, and 82; low-risk types: 6, 11, 30, 32, 34, 40, 42, 43, 44, 54, 55, 61, 62, 70, 72, 81, 83, 84, 90, and 91). DNA was extracted using a commercial kit (QIAamp Mini Kit, Qiagen, Valencia, CA, USA) on a robotic system, according to the manufacturer’s instructions. DNA was stored at – −70^0^ °C until use. HPV testing was undertaken first by PCR for amplification of a fragment of the L1 gene and human beta-actin, a housekeeping gene and reference gene.

DNA amplifications were performed using previously reported consensus primers. The sequence from 1024 to 1203 bp of the L1 gene (e.g HPV 16, GenBank No. K02718.1) was amplified using primer sets MY11 (5’-GCMCAGGGWCATAYAYAATGG-3’) and GP6-1 (5’-AATAAACTGTAAAT CATATTCCTC-3’). This region of HPV L1 is a highly conserved region despite integration of the HPV genome into the human genome and has been frequently analyzed in genotyping of HPV. The primer sets ACTBF (5’-GCACCACACCTTCTACAATGA-3’) and ACTBR (5’-GTCATCTTCTCGCGGTTGGC-3’) were designed to amplify the human beta-actin gene. The reaction mixture of 30 ㎕ contained 7 ㎕ of DNA sample, 23 ㎕ of PCR buffer (Perkin-Elmer, Norwalk, CT, USA), 3.5 mM MgCl2, 0.2 mM dNTP, 10 pmol of primer set of MY11/GP6-1, ACTBF/ACTBR, and 1 U of AmpliTaq polymerase (Perkin Elmer). The mixture was subjected to 40 cycles of amplification using a DNA thermal cycler 2720 (Perkin Elmer). For duplex PCR of the HPV L1 gene and human beta-actin gene, each cycle included a denaturation step at 95 °C for 30 s, an annealing step at 54 °C for 30 s, and a chain elongation step at 72 °C for 30 s. Each PCR was initiated with a 5-min denaturation step at 95 °C and finished by a 5-min extension step at 72 °C. To avoid false positives and false negatives, a negative control of reagent only (no template) and positive controls of recombinant plasmid carrying the L1 gene of HPV16 and HPV6 were included in each amplification reaction. PCR products were electrophoresed on 2% agarose gel (FMC Bioproducts, USA), stained with ethidium bromide, and photographed under UV light. PCR products of HPV L1 and human beta-actin gene were detected as fragments of approximately 180 bp and 102 bp, respectively, by electrophoresis. Specimens that tested positive by human beta-actin PCR (internal control), hence with adequate specimen quality, were subjected to HPV DNA genotyping.

#### HPV DNA microarray analysis

Sequences of the HPV L1 gene and human beta actin gene were amplified and labeled with Cy5. A mixture of 15 μL of amplified product and 50ul of distilled water was denatured by heating at 95 °C for 3 min, followed by cooling for 5 min on ice. The samples were mixed with 65 μL of GG hybridization buffer (Goodgene, Inc.) and then applied to the DNA chip. A perfusion 8 well chamber (Schleicher and Schuell BioScience, Germany) attached to the chip was used as a hybridization reaction chamber. Hybridization was performed at 50 °C for 30 min, followed by washing with washing buffer 1 for 2 min, washing buffer 2 for 2 min, and air drying at room temperature. The hybridization signal on the HPV DNA chip was visualized using a GenePix® 4000 B Microarray Scanner (Molecular Devices, Inc., Sunnyvale, CA, USA). The Ct value was obtained and analyzed for the degree of relative expression between the signal of the housekeeping gene and the signal for multiple HPV-related probes after the background noise was eliminated. The cutoff value for positivity was set at greater than 2.5.

The 40 types of HPV analyzed by this DNA microarray included 14 high-risk types of HPV, 6 probable high-risk HPVs, 13 low-risk HPVs, and 7 probable low-risk or undetermined-risk HPVs. The 14 types of HPV classified as high-risk include HPV type 16, 18, 31, 33, 35, 39, 45, 51, 52, 56, 58, 59, 68A/68B and 82. The six types of HPV classified as probable high-risk included HPV types 26, 53, 66, 67, 69, and 73. The 13 types of low-risk HPV include HPV type 6, 11, 34, 40, 42, 43, 44, 54, 55, 61, 62, 70 and 72. The seven probable low-risk or undetermined risk types of HPV included HPV30, 32, 73, 81, 84, 90, and 91. The classification of risk of various types of anogenital HPV in the current study was based on categorization of HPV types by expert working group at the International Agency for Research on Cancer (IARC) in 2009, epidemiological classification by the IARC Multicenter Cervical Cancer Study Group, and the naive Bayes classification. ^22–25^ We also referred to the classification as was used in the Hybrid Capture 2 Assay (Digene Corporation, Gaithersburg, MD, USA). Multiple HPV infections were defined as co-infection with two or more types of HPV.

#### HPV DNA sequencing assay

The genotyping data obtained from the HPV chip were further confirmed in 200 samples (188 specimens with mixed types on DNA chip + 12 specimens with ambiguous results close to Ct value cutoff on chip scanner) by conventional direct DNA sequencing method, as described previously, but with slight modifications. ^20^ In brief, 1-3ng/μl of the PCR amplifier of the HPV L1 gene was mixed with 8 μL of ABI Prism BigDye Terminator Cycle Sequencing Ready Reaction kit version 1.1(Perkin Elmer Biosystems, USA), 2 pmol of primer, and distilled water. The reaction mixture (10 μL) was treated for 5 s at 96°C, followed by 25 cycles of 96°C for 10 s, 50-C for 5 s, and 60-C for 4 min, followed by cleaning using a Centri-Sep 96 well plate or BigDye XTerminator Purification kit. After performing the post-sequencing reaction purification, samples were analyzed using an Applied Biosystems capillary electrophoresis-based genetic analyzer and then run on an ABI 3730xl instrument (Perkin Elmer, USA) and analyzed with DNA sequence data collection software (Perkin Elmer). The sequence data obtained by automated DNA sequencing were analyzed using a BLAST search (http://www.ncbi.nlm.nih.gov/BLAST/) for HPV genotyping.

## RESULTS

Figure 1 shows a representative view of genotyping results by DNA chip and DNA sequencing of various HPV types. Figure 2 shows a representative view of some examples of HPV results on sequencing and the HPV DNA chip. The results of the genotyping assay for 1,015 male genital warts are summarized in Table 1.

**Figure 1.**
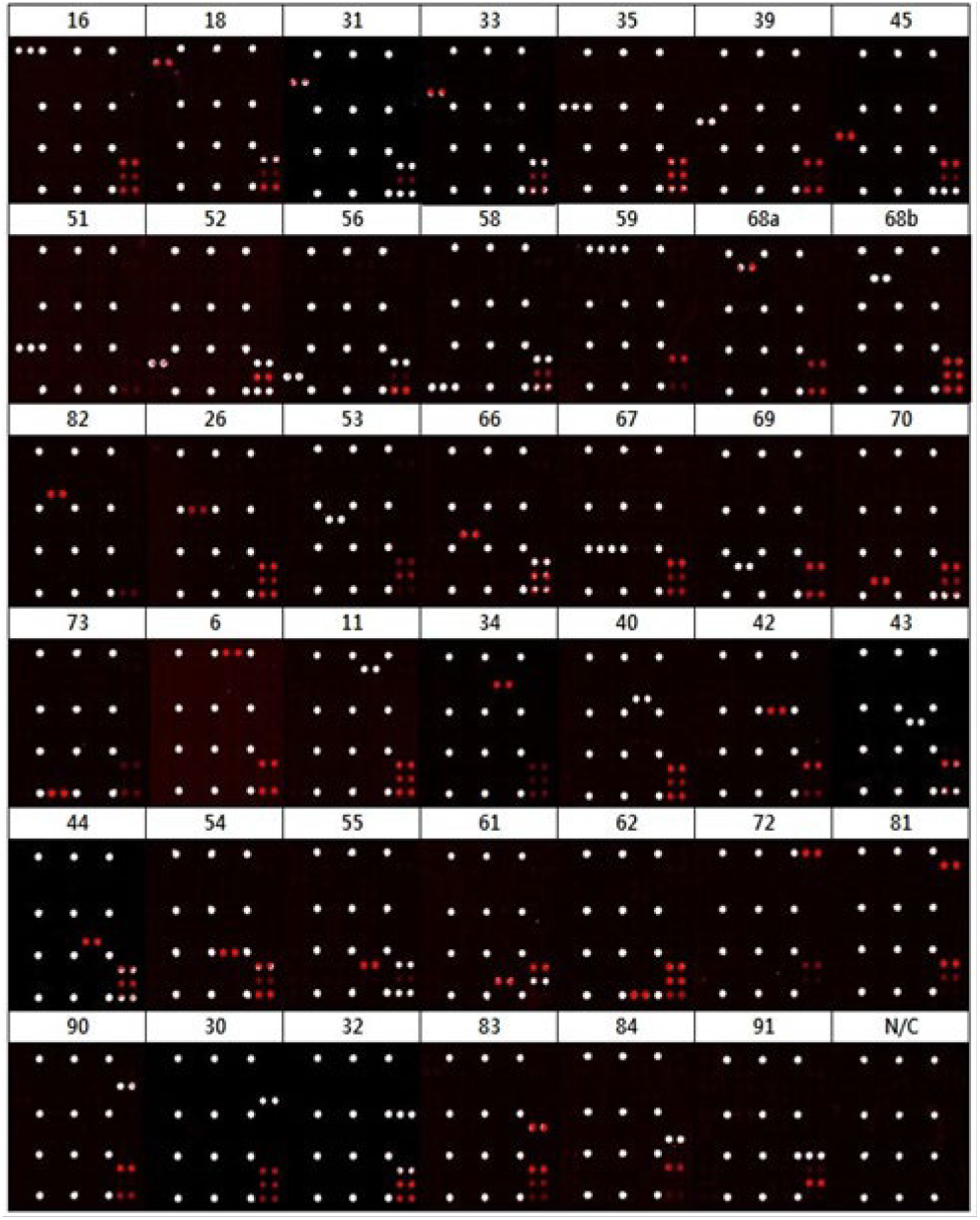
Representative view of HPV genotyping by using DNA microarray and sequencing assay

**Figure 2.**
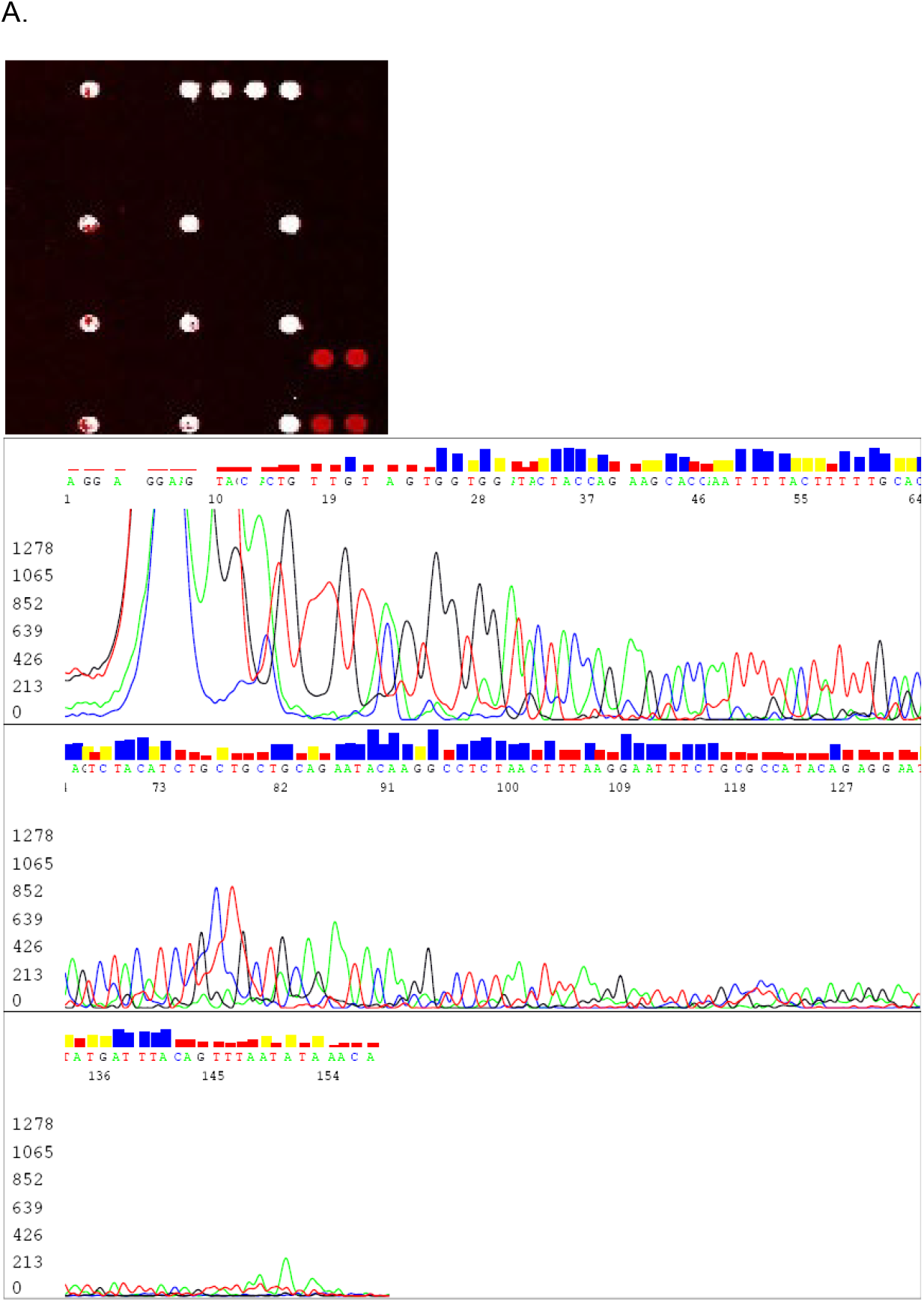
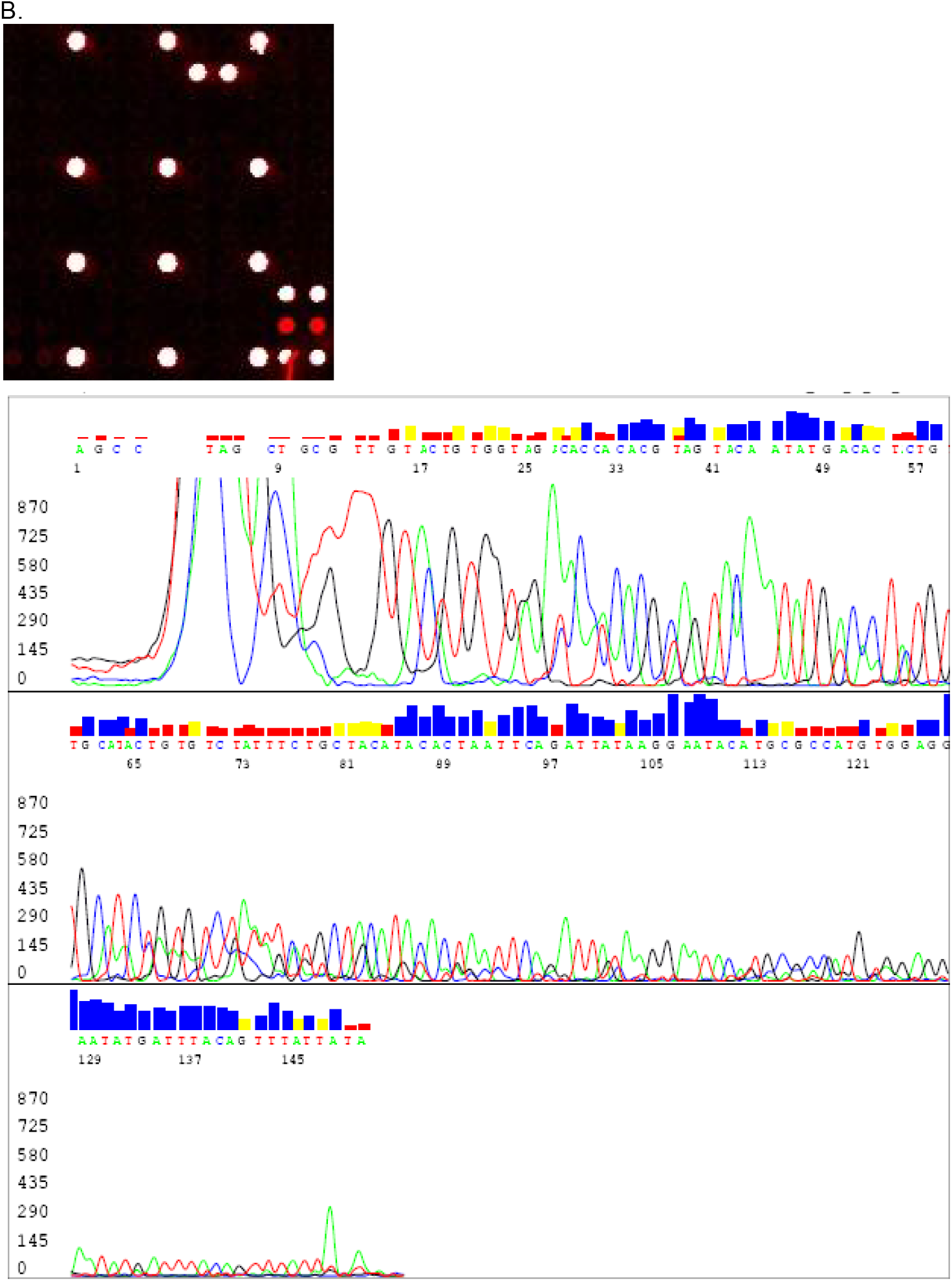
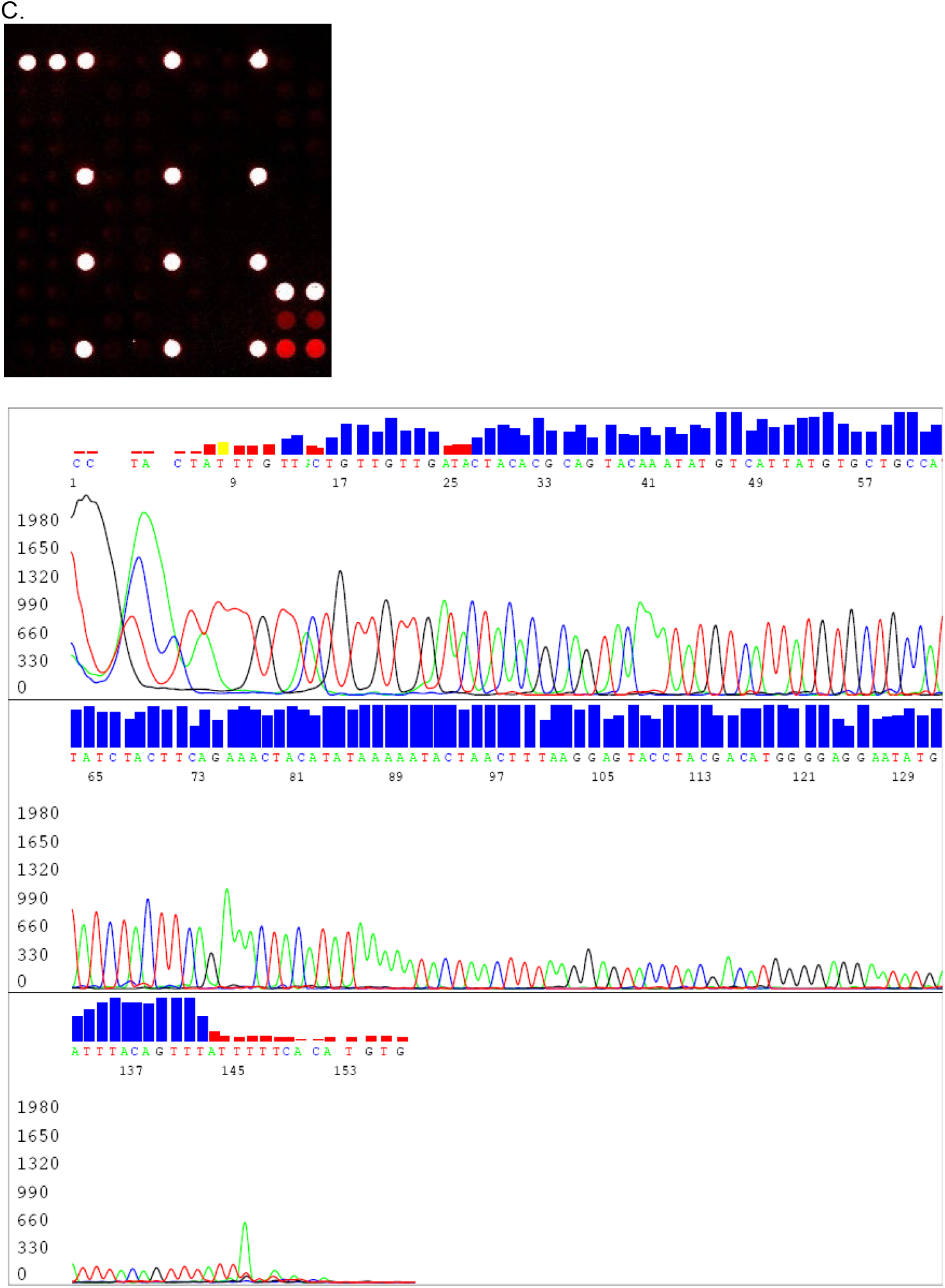
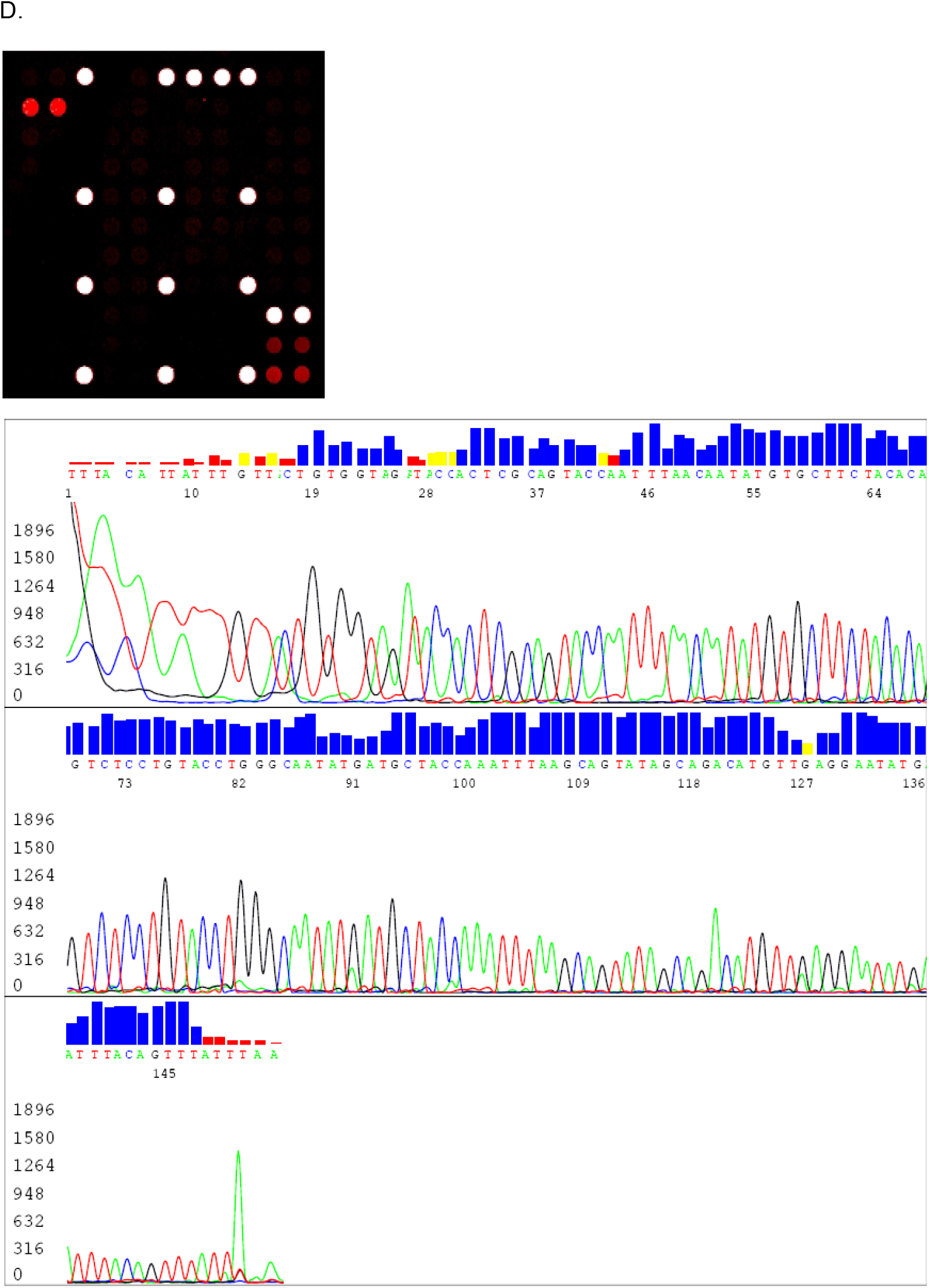
Genotypic distribution of 1,000 male genital warts as related with HPV6, HPV11, HPV16 and HPV18. A. HPV6 single infection B. HPV11 single infection C. HPV16 single infection D. HPV 6 and 18 mixed infection

**Table 1.**
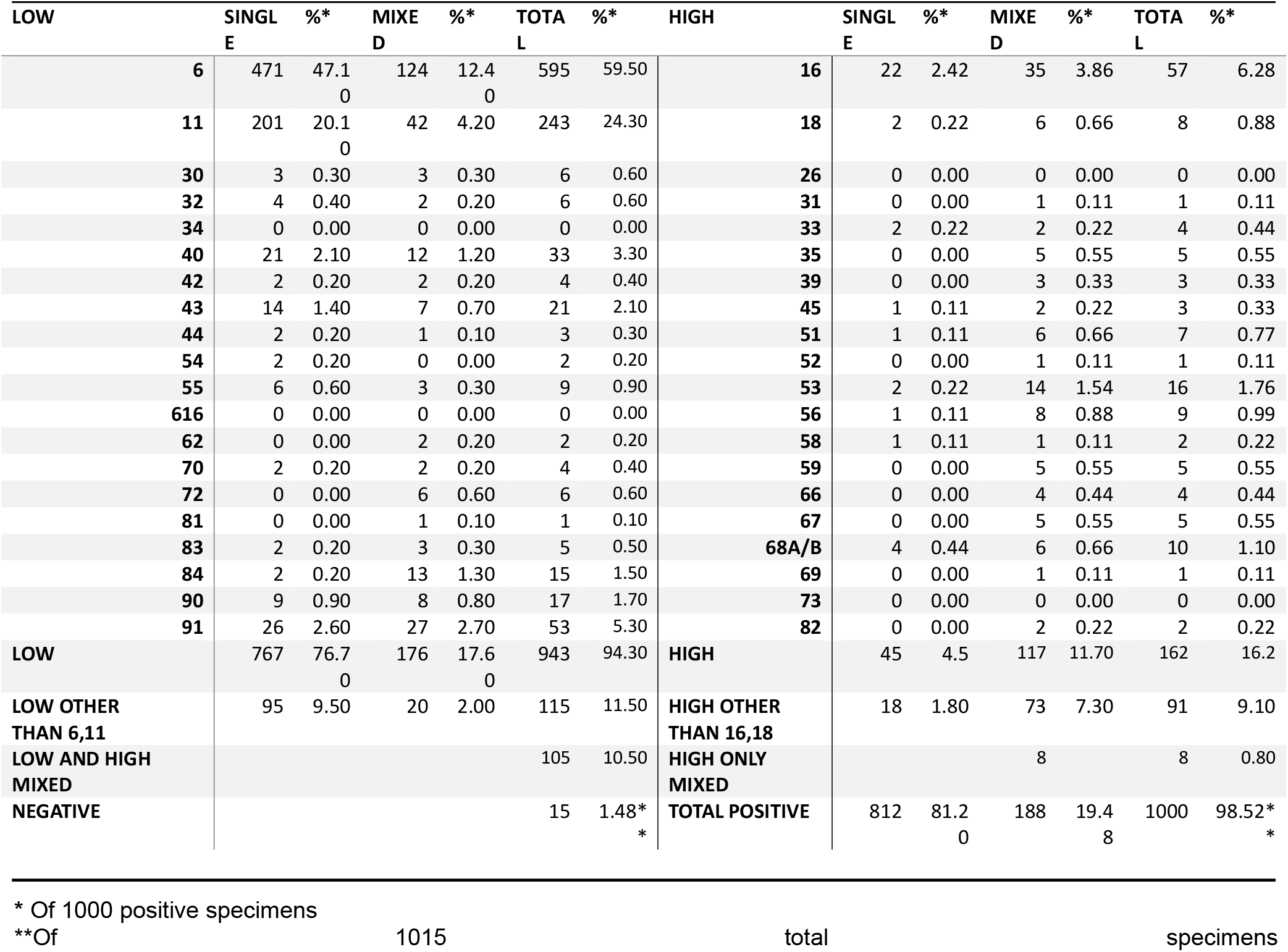
Genotypic distribution of HPV in 1,015 male genital warts.

Of the 1050 patients recruited, 35 patients were excluded from the study because of a lack of necessary information, improper sampling, or coexisting conditions. A total of 1,015 samples showed adequate quality, as defined by positive PCR results for the beta-globin gene. Out of these 1,015 adequate samples, 1000 (98.5%) showed HPV DNA after PCR on electrophoresis and hybridization on a DNA chip. Fifteen specimens did not show HPV DNA products after PCR by both electrophoresis and hybridization on the DNA chip and were excluded from the HPV genotyping analysis. In all 1000 HPV DNA-positive cases, HPV genotypic information could be obtained successfully by the HPV40 DNA chip assay. The age of the 1000 patients included in the final analysis ranged from 17 to 66 years, with an average age of 35 years. The majority of patients (85.8%) presented before the age of 40 years. The location of genital warts were as follows: penile shaft (35.2%), coronal sulcus (30.0%), base of penis (12.3%), glans (9.5%), scrotum (5.8%), perineum (4.3%) and perianal area (2.9%) in decreasing order of frequency.

Out of 1000 samples positive for HPV DNA on analysis by using DNA chip, 200 samples were selected with mixed infection (n=188) or equivocal (close to cutoff 2.5) findings on DNA chip (n=12) analysis. A DNA sequencing assay was also performed as a validation study to confirm the results of DNA chip analysis in the above 200 samples. The results of the genotyping study by DNA chip and DNA sequencing matched 184 samples. All the remaining 16 samples were found to have mixed infection due to more than one type of HPV, in which DNA sequencing detected only one or two types of HPV(s), whereas the DNA chip showed all types of HPV present in the sample. This was confirmed by a type-specific PCR assay. The results are summarized in Table 2.

**Table 2.** Sequencing Results vs HPV DNA chip results in 16 specimens with discrepancy. Samples were assayed for the appearance of HPV as described in the results section. ‘−‘ denotes not significantly positive during assay. All 16 samples were found to have mixed infection due to more than 1 type of HPV, in which DNA sequencing detected only 1 or 2 types of HPV(s), whereas DNA chip showed all the types of HPV present in the sample. HPV chip results were confirmed to be true by type specific PCR assay.

| SAMPLE NUMBER | HPV CHIP |  | SEQUENCING |  | CT VALUE (AVERAGE) |
| --- | --- | --- | --- | --- | --- |
|  | low risk | high risk | low risk | high risk |  |
| 7 | 6 | 53 | 6 | - | 2.9 |
| 40 | 6,91 | 18 | 6 | 18 | 2.7 |
| 90 | 6, 40 | 66 | 6 | 66 | 2.5 |
| 96 | 6, 40 | 16 | 40 | 16 | 2.8 |
| 128 | 6, 40 | 16 | 40 | 66 | 2.6 |
| 133 | 6, 84 | - | 6 | - | 3 |
| 237 | 6,11, 84 | - | 6, 11 | - | 3 |
| 359 | 11, 91 | 16 | 11 | 16 | 2.4 |
| 536 | 40, 69 | 16,18 | 40 | 16, 18 | 2.7 |
| 577 | 6, 72 | 16 | 6 | 16 | 2.5 |
| 640 | 40 | 16, 58 | 40 | 16 | 2.9 |
| 720 | 6 | 16,18 | 6 | 16 | 2.4 |
| 740 | 6 | 56, 82 | 6 | 82 | 3 |
| 767 | 6, 54 | 59 | 6 | 59 | 2.6 |
| 776 | 90 | 68, 82 | 90 | 68 | 2.7 |
| 906 | 6 | 35, 18 | 6 | 18 | 2.8 |

Depending on oncogenic risk, 94.3% of samples showed infection by low-risk HPV, 16.2% high-risk type HPV, and 10.5% both high-and low-risk HPV types, respectively.

Twelve types of low-risk HPV were found in the male genital warts. HPV 6 and HPV 11 were the two most common HPV types found, and they together accounted for 82.8% of all samples of male genital warts. 17.2% showed only HPV type(s) other than HPV6 and HPV11. HPV6 was found as single infection in 47.1% (471) and as mixed infection with other HPV type(s) in 12.4% (124) of positive samples. HPV11 was found as single infection in 20.1% (201) and as mixed infection with other HPV type(s) in 4.0% (40) of positive samples. However, HPV of types other than HPV6 and HPV11 were also found at high frequencies. HPV91, HPV40, and HPV43 were some of the frequent low-risk types found in our study.

Eighteen high-risk HPV types were found in the male genital warts. HPV 16 and HPV 18 together accounted for 82.8% of all male genital warts. HPV18 was found in only 16 cases (1.6%). High-risk HPVs other than HPV16 or 18 were found in low frequency (56.2% of high-risk / 91 total) and usually were found to be co-infected with low-risk HPV (n= 105, 89.7%).

Of the 188 samples that showed mixed HPV infection, the majority (167) presented with double infection, 20 with triple infection, and 1 with quadruple infection. The most common type was co-infection with HPV6 and HPV16 (n=21, 2.1%), followed by HPV 6 and 11 co-infection (n=10, 1%), and HPV6 and HPV 91 co-infection (n = n-10, 1%).

In the present study, the most common type of HPV was HPV6 (59.5%), followed by HPV11 (24.3%), HPV16 (6%), and HPV91 (5.3%). These results concur with the previous reports that HPV6 and HPV11 are the two most common HPV types found in male genital warts.^35^ However, in our study, they together accounted for not 90% but only ~82% of male genital warts. HPV of types other than HPV6 and HPV11 were also found at a high frequency (approximately about ~31.8%). 17.2% of genital wart samples showed neither HPV6 nor HPV11.

The most striking finding was that high-risk HPV, including HPV16, was found in 16.2% of male genital warts. HPV types other than HPV6, 11, 16, and 18, such as HPV91, HPV40, and HPV53, account for a considerable proportion (13.4%).

The distribution of HPV types was analyzed with respect to HPV6, HPV11, HPV16, and HPV18 as well as HPV31, 33, 45, 52, and 58 to investigate the potential protection of quadrivalent vaccines against genital warts that develop secondary to these four types of HPV (Figure 3). Of 1000 samples of male genital warts (86.6%), 866 showed HPV6, HPV11, HPV16, and HPV18. Of the genital wart samples, 87.2 % and 85.9% of all genital wart samples showed HPV6, 11, 16, 18, 31, 33, 45, 52, and 58.

**Figure 3.**
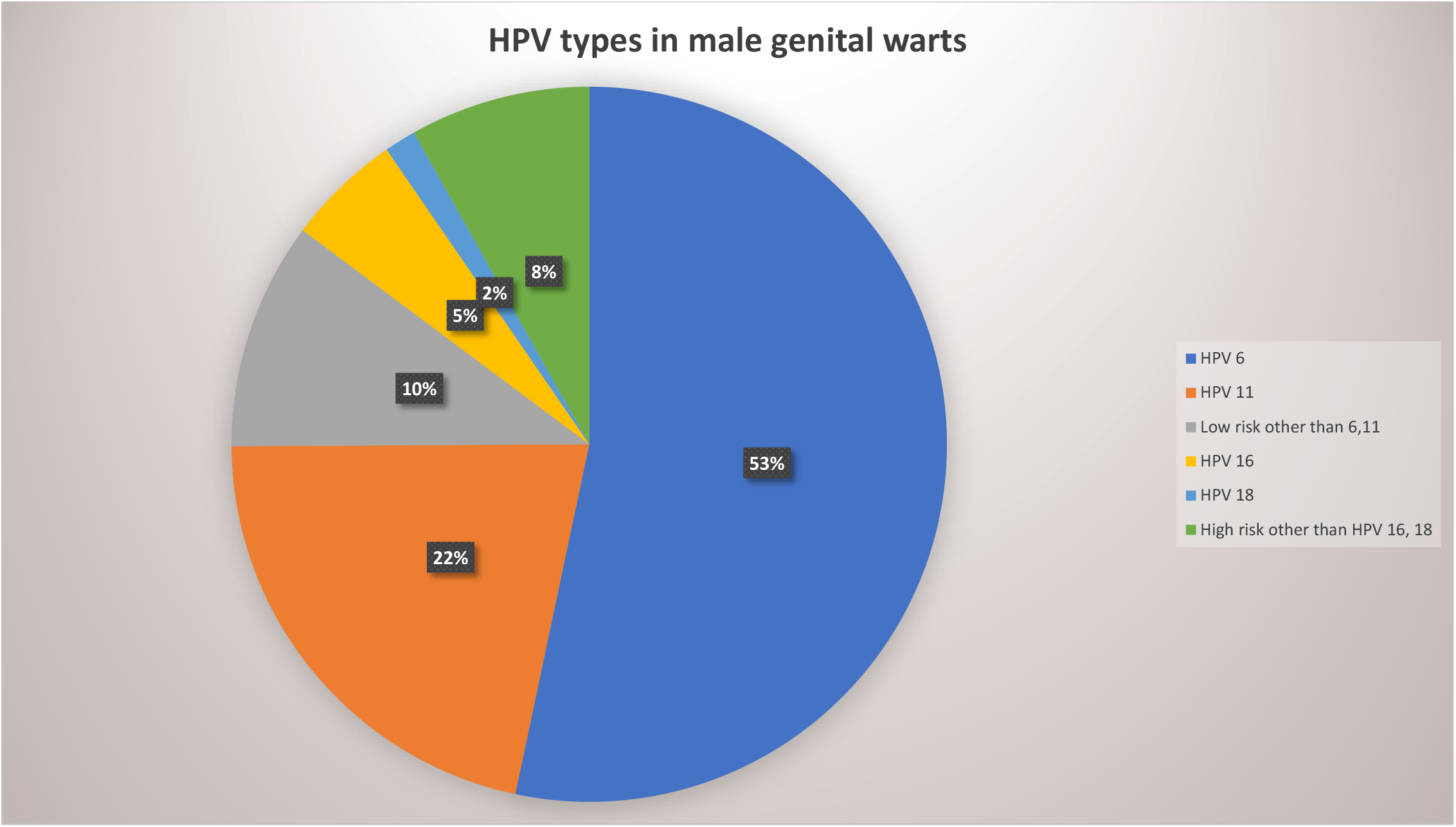
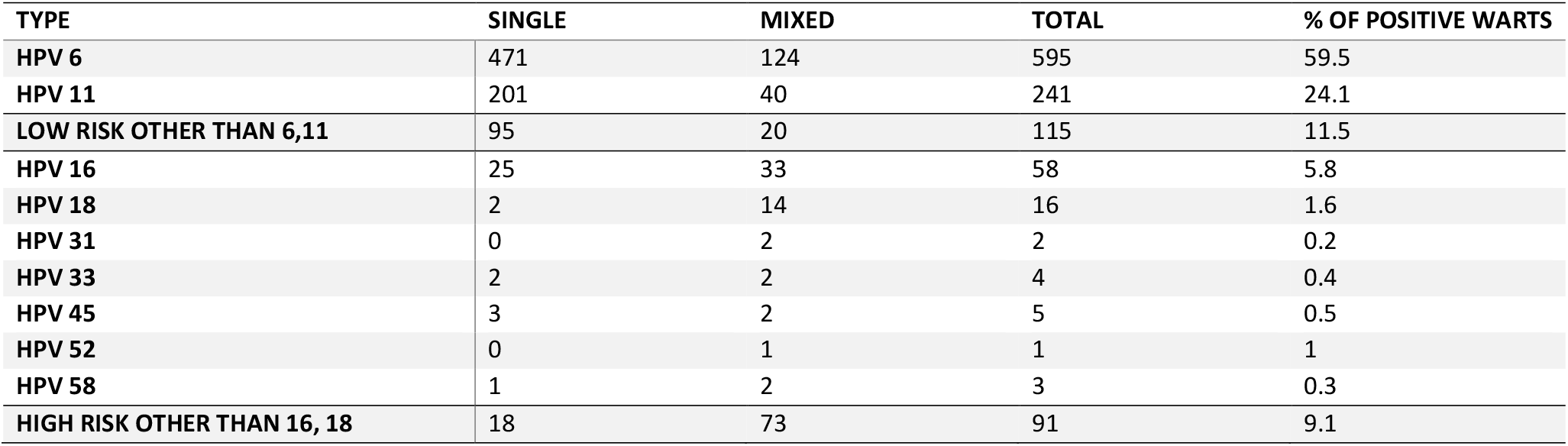
Genotypic distribution of 1,015 male genital warts as related with HPV6, HPV11, HPV16 and HPV18

## DISCUSSION

Genital warts are a significant public health problem. Genital warts are one of the most common sexually transmitted diseases (STDs), with an estimated 1–6 million new cases in the United States and about 30 million new cases worldwide each year.^14^ The National Survey of USA (1999–2004) in ~8,500 sexually active men and women reported an overall 5.6% history of genital warts. In addition, studies have shown increase in the occurrence of genital warts from 1966 to 2004 by 4-folds. Genital warts also incur significant healthcare costs for society. ^12–14, 27^

Although genital warts are known to be benign and not associated with mortality, they show a high recurrence rate despite apparently successful therapy. ^12, 13, 27^ They can also be rarely associated with malignancy in the form of Buschke-Löwenstein tumor, and warts associated with HPV 16 and 18 may be predisposed to oncogenic transformation. ^14,28^ In some studies, women with a history of genital warts have been shown to have an increased risk of CIN and cancer, which is most likely explained by a higher risk of having other oncogenic HPV types present. ^10, 29^ This observation is supported by nationwide studies that show that approximately 30%–40% of female genital warts contain oncogenic HPV type infection. ^19, 30^

There is increasing interest in understanding the burden of HPV infection and related diseases among men. Giuliano et al presented a prospective study on the incidence and clearance of HPV in men residing in Brazil, Mexico, and the USA. They were found to acquire a new genital HPV infection in high incidence (38.4 per 1000 person months). High-risk HPV types with the highest incidences were 16, 51, and 52, whereas low-risk HPV types with the highest incidences were 6, 62, and 84. About half of the men were found to harbor invisible HPV infection in the genitalia.^12^

There is also a lot of interest in the effects of HPV vaccines in men. Currently, the most recent prophylactic vaccine available for the prevention of HPV infection is Gardasil 9™ (Merck Sharp and Dohme) containing HPV 6, 11, 16, 18, 31, 33, 45, 52, and 58. The Gardasil 9 vaccine has been shown to have 97.4% efficacy in preventing high-grade cervical, vulvar, and vaginal disease for up to 6 years.^33^ While these vaccines primarily aim to prevent cervical cancers caused by high-risk HPV types, the inclusion of low-risk HPV types (HPV6 and 11) in Gardasil™ provides an additional benefit for preventing anogenital warts.^7,9,15,16,30^ The HPV vaccine is effective for the protection of anogenital diseases regardless of HPV type by 34%.^7,32^

Giuliano et al. reported the efficacy and safety of quadrivalent vaccine in the prevention of genital warts in heterosexual boys and men aged 16 to 23 years and male homosexual males aged 16 to 26 years, based upon which the United States FDA approved the use of the vaccine in boys and men aged 9 to 26 years for the prevention of genital warts.^7,8^ The Gardasil 9 vaccine also benefits public health by helping prevent HPV transmission to women as well as protection in homosexual men and is currently approved for individuals up to 45 years of age.^7,42^ In the per-protocol population, the efficacy of quadrivalent vaccine against lesions related to HPV-6, 11, 16, or 18 was 90.4%. However, in the intention-to-treat population, quadrivalent HPV vaccine showed an overall efficacy of 60.2%, and the efficacy was 65.5% for genital wart lesions related to HPV-6, 11, 16, or 18. These data suggest that vaccines show efficacy preferentially on genital warts related to HPV-6, 11, 16, or 18, and overall efficacy on entire genital warts is rather limited. It also suggests that up to 40% of genital warts develop due to HPV of types other than HPV6, 11, 16, and 18. ^15, 16, 32, 33,^

This study drew many interesting findings. Out of 1000 HPV-positive samples, a not-so-high proportion (18.8%) showed mixed infection due to more than one type of HPV. Our data from immunocompetent patients contained significantly lower rates of mixed infection than previous estimates (Brown et al., 1999; Aubin et al., 2008, Han et al., 2009). Siolian L. R. Ball has studied 31 wart tissues (both male and female, snapped frozen and immediately submitted for DNA extraction) that showed 71% mixed HPV infections and 48% high-risk HPV. Dual infections accounted for 35%, triple 10%, and equal to or more than quadruple (26%). This raises concern that in some of the cases in the present study, the sample collection and transport in the fresh tissue state in our study may have deteriorated some of the HPV DNA.

Overall, 32 types of HPV, including 18 types of high-risk HPV and 14 types of low-risk HPV, were found in male genital warts. These results suggest that contrary to common belief that HPV 6 and HPV 11 induce almost all genital warts in males, a variety of HPV types can be present in male anogenital warts, and high-risk HPV infection is more frequent than previously thought.

Ingles et al have conducted a similar study in which 77.2% of the EGLs were HPV positive with high-risk in 15.6% and low-risk types in 73.2%.^10^ In condylomas, 79.7% tested positive for low-risk HPV, 49.4% for HPV 6, 31.0% for HPV 11, and 8.2% for high-risk HPV, respectively.^10^ In lesions suggestive of condyloma, 75.8% tested positive for low-risk HPV, 57.3% for HPV 6, 18.5% for HPV 11, and 13.7% for high-risk HPV, respectively. Lesions with coinfections, including ≥1 high-risk and ≥1 low-risk HPV type, accounted for 6.3% and 12.1 %, respectively, which was in the range of the results shown in our study. HPV 6, 11, 16, 18, 31, 33, 45, 52, and 58 together accounted for 81.6% and 75.0%, respectively, similar to our study (85.9%).

These results indicate that currently available HPV vaccines targeting HPV6, 11, 16, 18, 31, 33, 45, 52, or 58. will be effective for the protection of male genital warts. According to previous studies, most 9vHPV vaccine recipients seroconvert for all 9 HPV types at month 7 and antibody responses to the nine HPV types persist over 5 years.^37^ It is suspected that the 9-valent vaccine may provide protection against >80% of male genital warts if we hypothesize that it can protect against 97% of genital lesions (efficacy calculated per Warner K Huh et al;^32^ ~97.4%/ Elmar A Joura et al;^33^ 96.7% in per-protocol population in cervical, vulvar, vaginal neoplasia) related to HPV6, 11, 16, 18, 31, 33, 45, 52, and 58 (prevalence in warts per our study: ~86%). This may be compared to Giuliano et al., who reported an efficacy of 90.4% for the quadrivalent human papillomavirus (HPV) vaccine against external genital lesions (mainly warts) related to HPV-6, 11, 16, or 18 in healthy, predominantly heterosexual males.^38^ In the intention-to-treat population, the observed efficacy of decrease in EGLs in the vaccine group is 60.2% and 65.5% for EGLs related to HPV-6, 11, 16, or 18.^38^ To protect against more than 90% of genital warts, a polyvalent vaccine that can include and protect against major types of HPV-inducing male genital warts, such as HPV91, HPV40, and HPV43 may be needed.

Given the high correlation of HIV with HPV, it may be suggested that HPV vaccination should be mandatory and anal wart/cancer surveillance is recommended in people with HIV. According to published meta-analysis, HPV incidence and high-risk HPV incidence approximately doubled among HIV patients and HPV clearance rate approximately halved. HIV incidence almost doubled in the presence of prevalent HPV infection.^39^ In another metanalysis, HIV-positive versus HIV-negative men had anal HPV-16 prevalence of 35·4% versus 12.5% and pooled anal cancer incidence was 45·9 per 100 000 men versus 5·1 per 100 000 men.^40, 41^

In addition, catch-up vaccination with booster shots and mixed vaccine schedules should be considered, especially in HIV-positive men. The study by Simoens et al. showed that gender-neutral vaccination (GNV) with catch-up 9vHPV vaccine showed reductions of 30.3% and 44.6% for genital warts in females and males, respectively.^36^ Studies have shown that HPV prevalence in MSM (men who have sex with men) is significantly higher than that in heterosexual men, whose HPV prevalence is similar to that of women,^41^ and HPV prevalence in HIV-positive men is significantly higher than that in HIV-negative men.^39–42^

The results of our study differs from that of previous studies in that we found that there were a lot of non-HPV 6/11 strains in the warts. ^4, 17–19^ The significance and role of high-risk HPVs in male genital warts, even though highly prevalent (16.2% in our study; 14%–44% in immunocompetent, 47%–100% in immunosuppressed and HIV-positive)^35^ is not clear. The detection of HPV DNA by DNA chip/sequencing alone does not directly indicate the presence of HPV infection. We cannot completely exclude the possibility of high-risk HPV DNAs being contaminated from other areas of genital areas or cancer/precancer tissue being included despite the pathologic diagnosis. A recent study on HPV in males showed a high prevalence of HPV (~50%) in grossly normal-looking areas of male genitalia, of which HPV 16 and other high-risk HPV were even higher than low-risk HPVs.^10^ However, high-risk HPV alone was found without co-infection with low-risk HPV in 53 samples (45 single, 8 mixed). A recent study indicated that precancerous lesions and even invasive cancers are found in high frequency within anal condyloma in MSM. This suggests caution for genital warts infected by HPV16 that may require meticulous follow-up of not only patients but also their sexual partners. Further studies on the significance and management of high-risk HPV DNA in genital warts are necessary.

The strengths of this study include the size of the cohort and the pathologic diagnosis of lesions to allow for accurate detection of lesions. To our knowledge, this is one of the largest scale HPV genotyping studies in male genital warts carried out in a prospective, multicenter study. In previous studies, up to a total of 755 patients were tested and they used a rather limited genotyping assay and tested HPV types of from 5 to 24 in number.^10, 17–19^ In the present study, we tested 1000 samples by using PCR followed by DNA microarray. GG HPV DNA microarray is a high-throughput assay that can detect the 40 types of HPV in a highly sensitive (detects 10-100 copies of HPV DNA) way. We also added a DNA sequencing assay in order to confirm and validate the results of the DNA chip.

Another strength includes the use of formalin-fixed biopsy tissue for genotyping, as the biopsy results have been shown to accurately reflect the HPV genotypes found within these lesions. In contrast to previous studies in which samples were taken from the surface of warts using a brush swab, which carries the risk of contamination from other areas of genital skin, we obtained fresh tissue fragments of warts by excisional biopsy in a sterile manner and tested its total genomic DNA. Most other studies have used PCR amplification with sequencing or hybridization that are so sensitive that they may also detect HPV DNA in the anogenital region in asymptomatic individuals.^35^

However, the present study has some limitations. The target population of the study was Korean men. Therefore, data from this study may not be directly applicable to other races or geographical areas. Further studies such as this may be mandatory in men from races other than Korean or Asian and areas in Western countries or Africa. However, the genotypic distribution of genital warts in the present study is similar to that of recent studies. Additional international studies are needed to understand the prevalence of HPV, especially high-risk HPV, in anal warts, and the impact of HPV vaccines on anal warts, especially in relation to HIV status and sexual orientation. The 15 HPV-negative wart samples and 16 samples with discrepant results between sequencing and DNA chip is also a problem. These might have not been real warts or warts but with HPV DNA copy of below detection limit.

In addition, the HIV positivity and sexual orientations of the study population have not been investigated; this should be a point of concern. Additional international studies are needed to understand the prevalence of HPV, especially high-risk HPV, in anal warts, and the impact of HPV vaccines on anal warts, especially in relation to HIV status and sexual orientation.

## Abbreviations and Acronyms

HPV: human papillomavirus
DNA microarray: DNA chip
IARC: International Agency for Research on Cancer
PCR: polymerase chain reaction
EGLs: External genital lesions
CA: Condyloma acuminatum
MSM: Men who have sex with men

## Author contributions

WM, JM, JJ, JY, and Korean Male HPV study group conceived of the presented idea. JJ and JY helped collect the samples and were involved in planning and supervising the work. WM and JM designed and performed the experiments, derived the models and analysed the data. WM and JM wrote the manuscript with support from JJ and JY.

